# Tissue-specific modulation of CRISPR activity by miRNA-sensing guide RNAs

**DOI:** 10.1101/2024.08.09.605335

**Authors:** Antonio Garcia-Guerra, Chaitra Sathyaprakash, Olivier G. de Jong, Wooi F. Lim, Pieter Vader, Samir El Andaloussi, Jonathan Bath, Jesus Reine, Yoshitsugu Aoki, Andrew J. Turberfield, Matthew J. A. Wood, Carlo Rinaldi

## Abstract

Nucleic acid nanostructures offer unique opportunities for biomedical applications due to their sequence-programmable structures and functions, which enable the design of complex responses to molecular cues. Control of the biological activity of therapeutic cargoes based on endogenous molecular signatures holds the potential to overcome major hurdles in translational research: cell specificity and off-target effects. Endogenous microRNAs can be used to profile cell type and cell state and are ideal inputs for RNA nanodevices. Here we present CRISPR MiRAGE (miRNA-activated genome editing), a tool comprising a dynamic single-guide RNA that senses miRNA complexed with Argonaute proteins and controls downstream CRISPR activity based on the detected miRNA signature. We study the operation of the miRNA-sensing single-guide RNA and attain muscle-specific activation of gene editing through CRISPR MiRAGE in models of Duchenne muscular dystrophy. By enabling RNA-controlled gene editing activity, this technology creates opportunities to advance tissue-specific CRISPR treatments for human diseases.

## MAIN

The Clustered Regularly Interspaced Short Palindromic Repeats (CRISPR)-Cas gene-editing technology utilizes RNA-guided nucleases (Cas proteins) to modify DNA or RNA duplexes at specific sites selected through their complementarity to a short guide sequence^1^. Due to its precision, ease of use, low cost and versatility, this technology has evolved rapidly for use in a vast range of applications including gene knock-out, gene knock-in, gene activation, gene repression, base editing and prime editing^2^. Myriad CRISPR-based clinical applications are expected to follow in the wake of recent successful human trials^3,4^. Nevertheless, off-target effects and genotoxicity have been shown to surge with increasing editing activity^5^ and still represent major obstacles to clinical implementation. Efforts to develop CRISPR systems with precise spatial and temporal control over expression and activity, employing a diverse set of genetic regulatory (e.g. cell-specific promoters)^6,7^, chemical (e.g. small-molecule activators and inhibitors)^8–10^, and physical (e.g. optical-, heat-, and ultrasound-responsive)^11,12^ approaches, have encountered significant challenges in translational research, due to factors such as insufficient tunability, system complexity, dependence to an exogenous activating stimulus, and background activity interference^5^. To this end, controlling CRISPR activity by engineering the guide RNA to respond to environmental cues (i.e. sensing endogenous molecules for tissue/cell specificity), capitalizing on research in the field of nucleic acid nanotechnology^13,14^, holds great promise. Integration of novel sensors that interact with molecular components of the cell to provide context-sensitive activation could introduce additional levels of control, enabling safer, more sophisticated CRISPR-based smart therapeutics. However, designing RNA nanodevices that fold into predictable and dynamic structures that interact predictably with endogenous molecules remains highly challenging^15^. Prior work mainly relies on miRNA-mediated guide RNA or Cas9 production^16^ or engineered guide RNAs that respond to a wide range of exogenous triggers, such as antisense oligonucleotides, small molecules, riboswitches and protein-coupled receptors^15,17–24^. We here present CRISPR MiRAGE (miRNA-activated genome editing), a technology whereas RNA devices, integrated in the CRISPR guide strand, change state to control the activity of CRISPR associated protein 9 from *Streptococcus pyogenes* (SpCas9) upon detection of a specific endogenous microRNA. MiRNAs are small, non-coding RNAs, present both in cytosol and nucleus, whose critical functions include transcript regulation and whose expression profiles are characteristic of tissue identity and disease state^25^. They carry out their modulatory functions upon association with Argonaute proteins (AGO) and are essential components of the RNA-Induced Silencing Complex (RISC)^26^. The RISC complex mediates gene silencing through transcript cleavage or translational repression based on its protein composition and the degree of target complementarity of the miRNA guide sequence^26^. AGO organises the miRNA into four functionally distinct domains^26^: seed region (nt 2-8), central region (nt 9-12), 3’ supplementary region (nt 13-16), and tail (nt 17-21). The seed is essential for miRNA binding and even targets with imperfect sequence complementarity are amenable to miRNA recognition. To find targets, AGO scans mRNA molecules using nucleotides 2-4 and, when there is a positive match, changes conformation to allow full hybridization of the seed^27^. The seed sequence of the RNA is arranged within AGO in such way to reduce energy penalties during hybridization^28^, with thermodynamic and kinetic properties typical of an RNA-binding protein. Our design and development of the conformation-switching CRISPR MiRAGE RNA guides, described below, is informed by the hypothesis, supported by experimental results, that their interaction with triggering miRNAs is mediated by AGO-miRNA complexes rather than simple strand displacement.

To explore the potential of CRISPR MiRAGE to increase the spatial and temporal precision of CRISPR therapeutics we use a fluorescent reporter system to demonstrate miRNA-specific activation of gene editing. We also validate tissue-specific CRISPR MiRAGE in *in vitro* and *in vivo* models of Duchenne muscular dystrophy (DMD), a fatal muscular disease that primarily affects skeletal muscle. DMD is caused by mutations in the *DMD* gene and is at the forefront of gene editing development^29^, illustrating the potential of this technology for tissue-restricted gene editing applications.

## RESULTS

### Design of miRNA-sensing sgRNAs

CRISPR gene editing is typically controlled by a single guide RNA (sgRNA)^30^ comprising an RNA molecule containing a transposable ∼20 nucleotide guide sequence complementary to the target DNA (crRNA) and a trans-activating CRISPR RNA (tracrRNA) domain with an essential secondary structure. Disrupting these elements prevents Cas9 from carrying out its gene editing function^31^. The core design of our miRNA-sensing sgRNAs relies on a self-complementary “trigger” hairpin that sequesters the guide sequence in the crRNA region, disabling Cas9 activity unless the hairpin is disrupted by a miRNA-induced conformational change mediated by AGO (OFF-to-ON state). Specifically, we introduced upstream of the guide sequence a miRNA binding site followed by a “shield” sequence that completes the trigger hairpin (**Fig. 1a**). AGO-mediated binding of the activating miRNA to its complementary binding site destabilizes the hairpin, resulting in the displacement of the competing shield sequence. Sequences of these functional modules, and of all complete MiRAGE guides tested, are recorded in **Supplementary Table 1**.

**Fig. 1:**
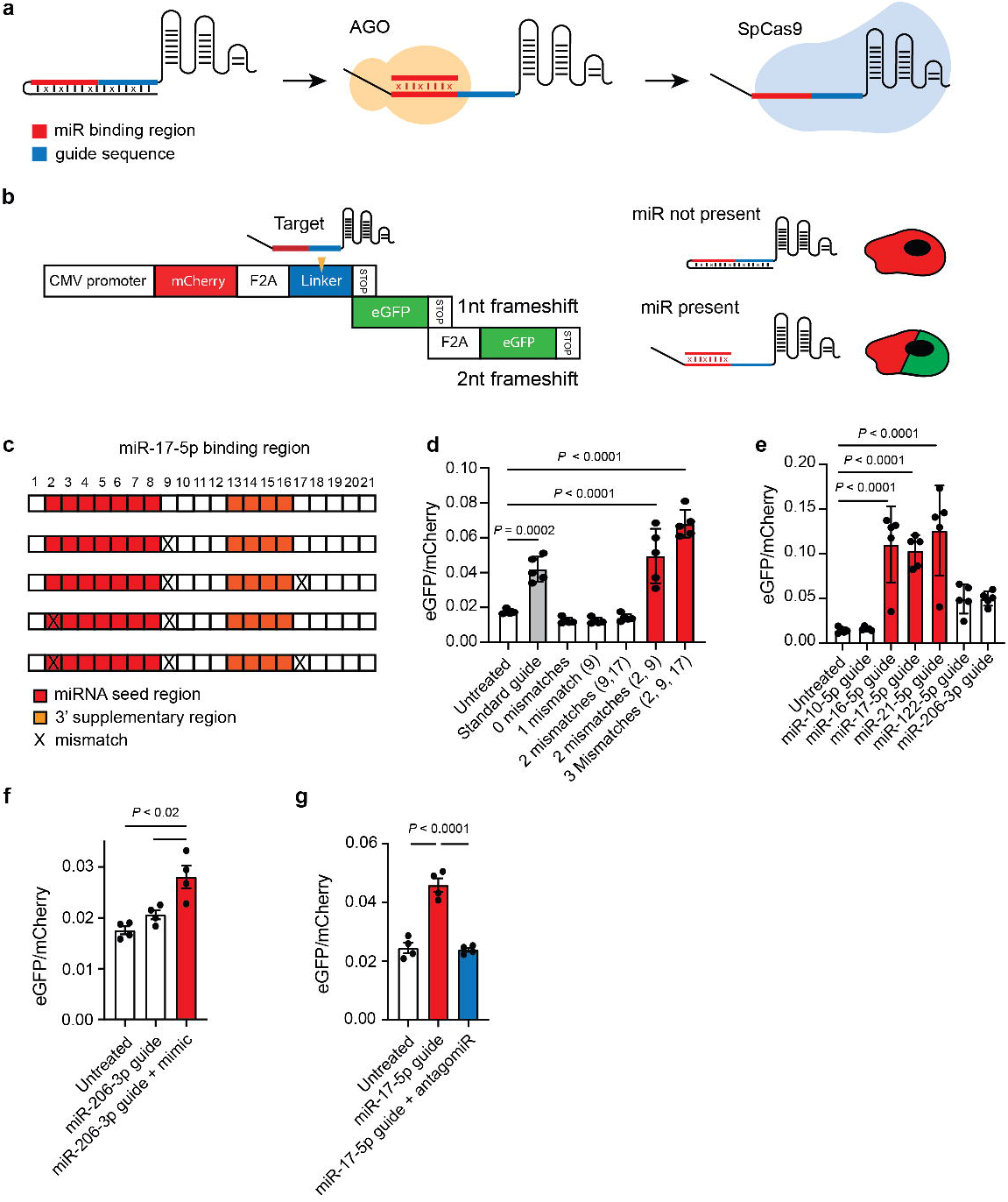
CRISPR MiRAGE design and activity. **a**. Schematic of the miRNA-responsive miR-guide design and its proposed operating mechanism. **b**. Structure of the Stoplight reporter construct. A constitutively expressed mCherry fluorescent protein open reading frame (ORF) is followed by two out-of-frame eGFP ORFs. eGFP expression results from introduction of insertions/deletions (INDELS) in the linker region through SpCas9-mediated double strand breaks. **c**. Different binding sites for miR-17-5p used in the sgRNA and shown in **d**. Mismatches were introduced in positions 2, 9, and 17 of the miRNA binding site, indicated with an X. **d**. Activities of miR-17-5p sgRNAs in the HEK293T Stoplight reporter cells. Positions of the mismatches are indicated within parentheses (*n* = 5). **e**. Performance of miR-guides containing mismatches in positions 2, 9, and 17, and responsive to a series of miRNAs (miR-10a-5p, miR-16-5p, miR-17-5p, miR-21-5p, miR-122-5p, and miR-206-3p). The activities of the miR-guides mirror the miRNA activity profiles measured in Stoplight cells (Supplementary Fig. 1c) (*n* = 5). **f**. miR-206-3p rescue experiment using miR-206-3p mimics. A miR-206-3p miR-guide is activated after co-transfecting exogenous miR-206 mimics (*n* = 4) **g**. The activity of miR-17-5p is prevented by treating the HEK293T Stoplight cells with an antagomiR (Cf = 10 nM) (coloured in blue) (*n* = 4). **d-g**. All data were analysed using one-way ANOVA and Dunnet’s multiple comparison test, except in **g** which used Tukey’s multiple comparison correction. The data represent the mean ± S.D. Histograms for standard guide and significant miR-guides are coloured in grey and red, respectively.

The intrinsic stability of the trigger hairpin is an essential feature: if too stable, AGO-mediated miRNA binding is inhibited, and if too labile, the ability of the guide strand to initiate Cas9 editing is insufficiently suppressed^32^. We used NUPACK^33^, based on the nearest-neighbour model of RNA thermodynamics^34^, to calculate the equilibrium ensemble of secondary structures. We tuned the stability of the hairpin by introducing mismatched nucleotides, distributed throughout the shield domain, to ensure that each base pair had a probability of approximately 90% of being hybridized at equilibrium as a compromise between sensitivity to the activating miRNA signal and a leak-free OFF state.

To assess the activity of our miRNA-sensing sgRNA (miR-guide) we used the Stoplight reporter system^35^ (**Fig. 1b**), which constitutively expresses fluorescent proteins mCherry and two out-of-frame eGFPs downstream from a linker region targeted by CRISPR, also SpCas9 (HEK293T Stoplight^+^ SpCas9^+^). Two thirds of non-homologous end joining (NHEJ) repair events within the linker region result in insertions-deletions (INDELs) that restore the reading frame of one of the eGFP genes, allowing direct visualization of successful editing by measurement of eGFP fluorescence.

To inform the choice of miRNA with which to test our miRNA-responsive guide technology, we developed a system for efficient functional screening of miRNAs in mammalian cells. We generated a second dual-luciferase reporter system in which a miRNA binding site is introduced downstream from a firefly luciferase gene. Upon transfection in HEK293T Stoplight cells, constructs containing binding sites for miRNAs that are not active in these cells (miR-122-5p and miR-206-3p) showed no activity (change in firefly luciferase expression), as expected. However, while constructs responsive to some of the highly-expressed miRNAs (miR-16-5p, miR-17-5p, and miR-21-5p) showed high activity, no signal was detected from a construct responsive to miR-10a-5p, despite high expression of the corresponding miRNA (**Supplementary Fig. 1a-c**). This corroborates previous evidence that miRNA expression and activity do not necessarily correlate in mammalian systems^36^ (**Supplementary Fig. 1c**,**d**). We chose miR-17-5p as an initial trigger with which to develop the CRISPR MiRAGE technology.

Upon transfection of our first-generation miR-17-5p-responsive sgRNA (miR-17-5p guide) in the Stoplight cells we observed no evidence of gene editing (**Fig. 1d**). Motivated by our hypothesis that AGO plays a key role in miRNA target binding and that the secondary structure of the substrate duplex (formed between the miRNA and its binding site) affects AGO activity^28,37^, we produced a series of miR-guides containing different patterns of mismatches within the miRNA binding site that are known to increase AGO turnover (**Fig. 1c**)^26,28^. We observed a significant increase of eGFP signal, indicating successful editing to a degree comparable or superior to a standard sgRNA, when mismatches were introduced in positions 2, 9, and 17 of the binding site. Notably, modification site at position 2 lies within the miRNA seed region (**Fig. 1d**). These results suggest that, although AGO-mediated miRNA binding is essential for activation of our trigger construct, too-stable binding by AGO inhibits Cas9 activity. To better understand the productive interaction between AGO and our miR-guide, we introduced a set of mismatches (in positions 10 and 11) known to abolish AGO cleavage^26^. The corresponding miR-guides successfully induced gene editing (**Supplementary Fig. 2a**,**b)**, suggesting that AGO binding to the trigger hairpin is enough to promote miR-guide activation. This is consistent with the observation that mammalian AGO preferentially represses its targets rather than cleaving them^26^. Elucidation of the interplay between AGO and Cas9 through their mutual interaction with the miR-guide requires further research.

In order to further validate our design, we confirmed that our rationally designed sgRNAs were miR-specific (**Fig. 1e**) and that gene editing activity mirrored the miRNA functional profile (**Supplementary Fig. 1c**). To further confirm the specificity of miRNA control of CRISPR MiRAGE, we showed that editing activity in the Stoplight cells could be induced upon exogenous transfection of a miRNA not present in these cells (miR-206-3p) (**Fig. 1f**) and prevented upon repression of the corresponding endogenous miRNA (miR-17-5p) (**Fig. 1g** and **Supplementary Fig. 1d**,**e**). We also confirmed that the miR-guides did not affect mRNA homeostasis: levels of known mRNA targets regulated by miR-17-5p (*ATG2B, NUP35*, and *TMEM127*) were unchanged upon expression of a miRNA 17-5p-sensing sgRNA, compared to both a standard guide and a miR-guide responsive to other miRNA (miR-16-5p guide) (**Supplementary Fig. 3**). These results demonstrate the sgRNAs can be rationally designed to modulate CRISPR editing activity based on a miRNA signature.

### Characterization and optimization of the CRISPR MiRAGE mechanism

Structural predictions for the optimized miR-17-5p guide by NUPACK^33^, which calculates the equilibrium distribution of secondary structures, and by CoFold^38^ which takes into account co-transcriptional folding, are identical (**Supplementary Fig. 4a-c)**, indicating that the secondary structures of miR-guides are independent of folding pathway^39^. NUPACK predicts that the stability of the self-complementary trigger domain is such that no significant strand displacement is induced by free miR-17-5p (**Supplementary Fig. 4d**), consistent with our hypothesis that the conformational change that underlies the observed miRNA-specific activation of the engineered guide strand is triggered by AGO-mediated binding rather than simple strand displacement^40,41^.

To study the impact of structural changes within the trigger hairpin, we increased the number of base pairs formed within the trigger hairpin, while keeping the miRNA binding site constant correspondingly lengthening the shield sequence. This results in the creation of miR-17-5p guides with trigger hairpins with increasing stabilities (ΔG = −15, −19, −25, and −30 kcal/mol). We observed an inverse correlation between editing performance and stability (**Fig. 2a,b**), consistent with the prediction that AGO-mediated miRNA binding is limited by the stability of the competing trigger hairpin^37^. We also explored the effect of increasing the number of uninterrupted base pairs that the shield domain forms in the seed region of the guide sequence (positions 1-10 upstream from the PAM)^42^ while introducing compensating mismatches elsewhere to keep the free energy of the trigger hairpin approximately constant (ΔG = ∼ −15 kcal/mol): this strongly decreased editing activity (**Fig. 2c,d**). We conclude that the stability of the trigger hairpin, which can be tuned through the introduction of mismatches in the shield sequence, is a key parameter, and that the distribution of mismatches within and between the miRNA-binding and Cas9 guide sites is also important.

**Fig. 2:**
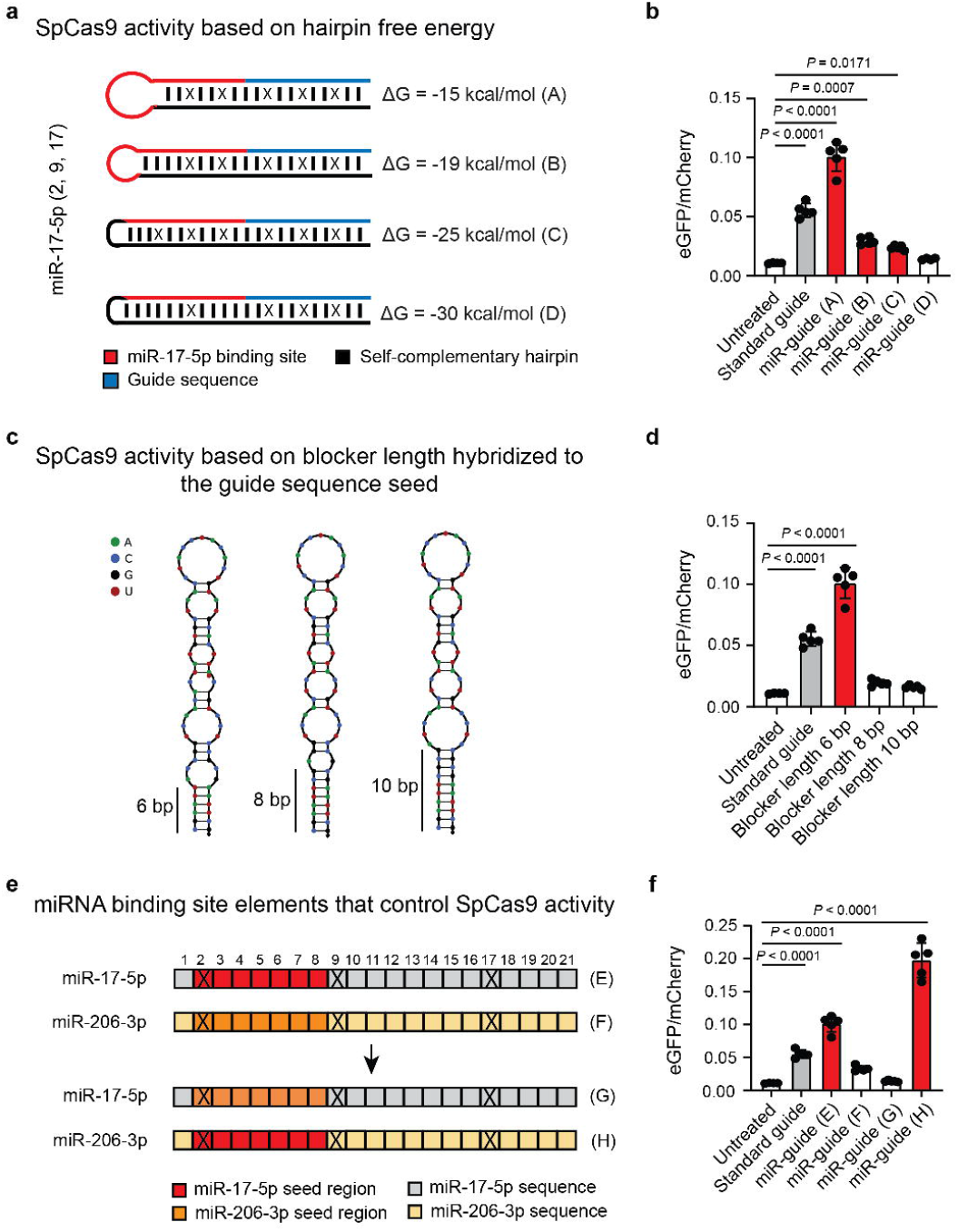
RNA secondary structure affecting CRISPR-MiRAGE. **a** Diagram depicting the different sgRNAs designed to test the impact of hairpin free energy on CRISPR-MiRAGE performance. While the trigger hairpin design is the same, the shield sequence is modified to form more base pairs and increase stability. **b** Impact of variations in free energy on miR-guide performance. Under tested conditions, eGFP signal decreases as hairpin stability increases, (*n* = 5). **c**. Diagram of miR-guide hairpins with different lengths of continuous duplex (6, 8 and 10 bp) at the guide seed sequence. Each hairpin contains the same number of mismatches to keep a constant free energy (−15 kcal/mol). **d**. Effect on miR-guide activation of the number of continuous bases hybridized to the guide seed sequence. Only the version with 6 base pairs blocking the guide sequence seed is capable of eliciting significant activation (*n* = 5). **e**. Diagram showing miR-guides with transposed seed sequences used to identify the essential elements of the miRNA binding site. Positions of the mismatches are indicated with an X. **f**. Seed swapping experiment. The inactive miRNA 206-3p sgRNA becomes active when the seed region is swapped with miR-17-5p seed region. Conversely, the miR-17-5p sgRNA is inactivated upon swapping with miR-206-3p seed region (*n* = 5). **b**,**d**,**f**. All data were analysed using one-way ANOVA and Dunnet’s multiple comparison test. The data represent the mean ± S.D. Histograms for standard guide and significant miR-guides are coloured in grey and red, respectively.

As the seed region of a miRNA provides most of the energy for AGO-mediated binding to the miRNA’s target^28^, we tested the role of the miRNA seed in activating our miR-guides. We introduced the miRNA seed region of an inactive (not expressed) miRNA (miR-206-3p) within the miRNA binding region of an active miRNA (miR-17-5p) and *vice versa*, and observed that editing activity mainly relies on specificity of the miRNA seed region (**Fig. 2e,f**). MiR-guides containing only the miRNA seed region and a mismatch in position 2 showed the same efficacy as their counterparts containing a full-length miRNA binding site (**Fig. 3a,b**). We next sought to test the impact of the position of the trigger hairpin within the guide RNA: accordingly, we developed an alternative design where the shield sequence is moved to the tetraloop region within stem-loop 1 (**Supplementary Fig. 5a)**. This alternative miR-guide successfully edits HEK293T Stoplight^+^SpCas9^+^ (**Supplementary Fig. 5b)**. We confirmed that predictions of the structural features of the alternative miR-guide are identical regardless of folding pathway (**Supplementary Fig. 5c**,**d)**. Overall this data reinforces that AGO interactions set the design rules for our miR-guides.

**Fig. 3:**
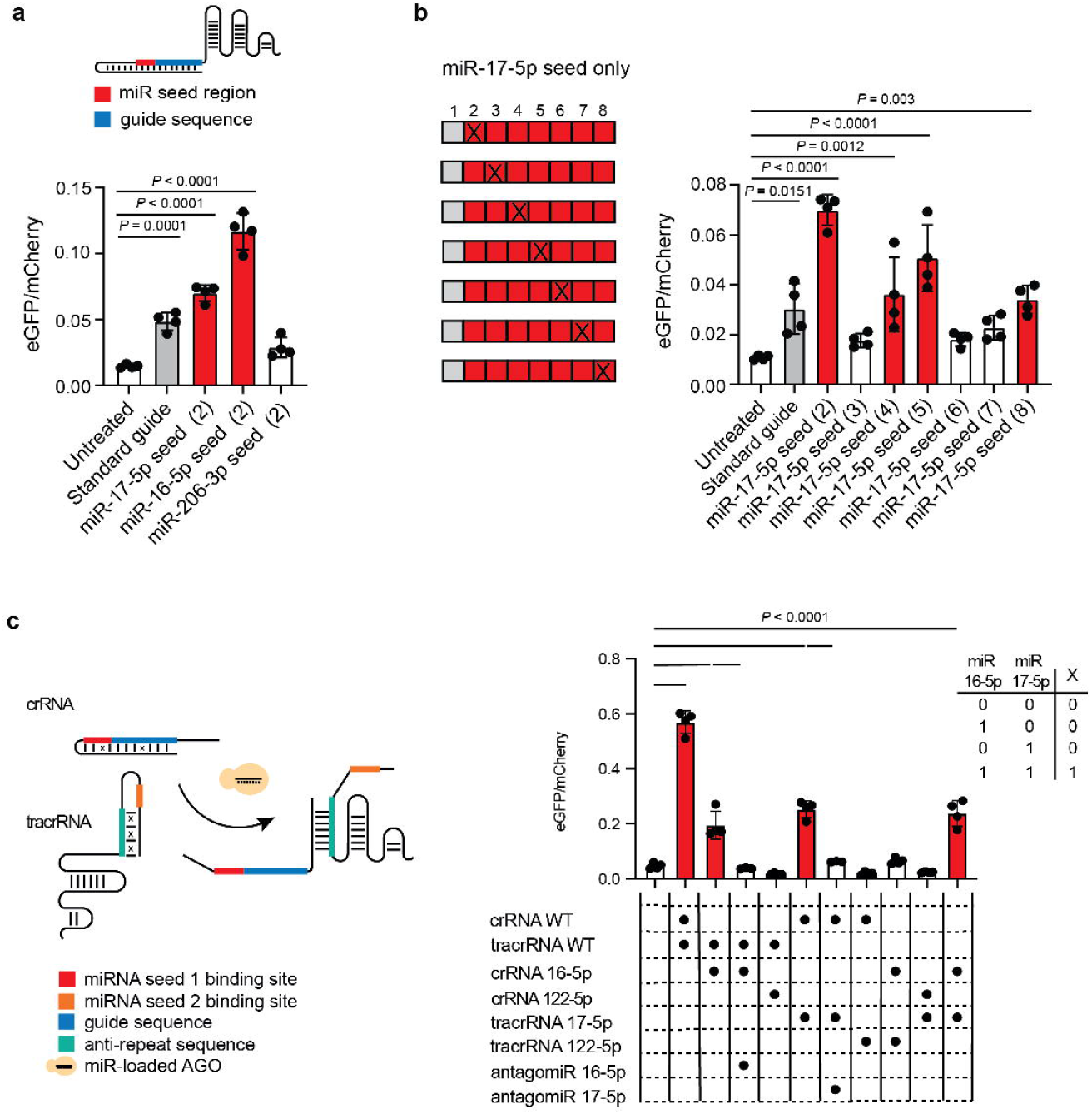
CRISPR-MiRAGE optimization. **a**. Top, a diagram showing a miR-guide where only the target miRNA seed is included. Bottom, activity, expressed as eGFP/mCherry ratio, for the seed-only miR-guides (*n* = 4). **b**. The impact of a single mismatch, indicated with an X, was tested as a function of its position across the miRNA seed. We observed a 3-base periodic trend which may reflect a strongly position-dependent destabilising effect of the mismatch on the RNA duplex or on the RNA-AGO interaction^26,57^ (*n* = 4). **c**. Left, a split miR-guide design results in multi-input-dependent activity. Right, performance of an AND-gated miR-guide in presence of active miRNAs (miR-16-5p and miR-17-5p), their relative antagomiRs, and inactive miRNA (miR-122-5p) are shown. This split design system follows a Boolean AND gate behaviour, as shown in the top right AND truth table. **a**,**b**. All data were analysed using one-way ANOVA and Dunnet’s multiple comparison test, except in **c**. which used Tukey’s multiple comparison correction. The data represent the mean ± S.D. Histograms for standard guide and significant miR-guides are coloured in grey and red, respectively.

We proceeded to optimize the activity of the guide by altering structural and sequence motifs within the sgRNA backbone. The activity of our CRISPR MiRAGE was improved by >75-fold when previously described backbone modifications (i.e. extending the hairpin derived from the native crRNA-tracRNA duplex and mutating one nucleotide of a continuous stretch of uracils to a cytosine) were introduced to increase Pol III transcription efficiency^43^ (**Supplementary Fig. 6a**), displaying a direct dose-responsive effect (**Supplementary Fig. 6b**). We further validated the miRNA-specificity of our optimized miR-guides by testing 4 more miRNAs against their respective antagomiR (i.e. miR-18a-5p, miR-20a-5p, miR-106a-5p, let-7a-5p) (**Supplementary Fig. 6c and Supplementary Fig. 1d)**. Additionally, we confirmed the activity of a model miR-guide (i.e miR 18a-5p) using flow cytometry (**Supplementary Fig. 6d-f)**.

Having characterized and optimized miR-guides transcribed with the cell, we confirmed that guides introduced by transfection following *in vitro* transcription were also active (**Supplementary Fig. 6g)**. CRISPR MiRAGE sgRNAs could therefore be delivered for clinical applications either via viral delivery or as chemically synthesized RNA payloads via, e.g., lipid nanoparticle technologies^44,45^.

In order to extend the translational potential of CRISPR MiRAGE and in consideration of the fact that a set of miRNAs may constitute a more specific signature of tissue type and/or disease state than a single species, we implemented a multi-miRNA-sensing version of CRISPR MiRAGE. We designed a two-part guide in the optimized backbone (**Supplementary Fig. 6a**), containing trigger hairpins blocking the guide sequence within the crRNA and the anti-repeat region within the tracrRNA, each of which is opened in response to a different miRNA (**Fig. 3c**). This split system requires both inputs (i.e. miRNA 17-5p and miRNA 16-5p) for activation of editing (**Fig. 3c**).

### Testing CRISPR MiRAGE in models of Duchenne muscular dystrophy

In order to validate this technology in mammalian systems, we used models of Duchenne muscular dystrophy (DMD) to test, as proof of principle, the ability of CRISPR MiRAGE to restore protein expression in a tissue-restricted manner. DMD is a severe muscular wasting condition caused by loss-of-function mutations in the *DMD* gene at the forefront of therapeutic gene editing development^29^. To this end, we used immortalized myoblast cells from DMD patients carrying a deletion in exon 52 (Δ52) which, by producing a premature stop codon in exon 53, results in loss of dystrophin expression (**Fig. 4a**). To achieve muscle-specific gene editing, we employed an optimized miR-guide with a validated guide sequence targeting the exon 53 splicing acceptor site^46^ and containing as trigger sequence the seed-binding domain for either muscle-specific miR-206-3p^25^ (myo-miR-guide) or, as a control, a liver-specific microRNA which is not expressed in skeletal muscle, miR-122-5p^25^ (liver-miR-guide). Notably, miR 206-3p shares the same seed sequence as miR 1a-3p, a miRNA highly expressed in cardiac muscle^49^ which is also severely affected in this disease. Upon successful gene editing, dystrophin expression can be restored by either exon skipping or exon reframing (**Fig. 4a**). Electroporation of the Δ52 myoblasts with a plasmid encoding SpCas9 and a miR-guide with the anti-dystrophin guide sequence showed that the myo-miR-guide and not the liver-miR-guide produced successful editing, as measured by trace analysis using Deconvolution of Complex DNA Repair (DECODR)^50^ of dystrophin amplicons (although to a lesser degree than the standard sgRNA (Std. guide) targeting the Δ52 mutation - positive control)^46^ (**Fig. 4b**). Analysis of the editing profile was consistent with single-cut activity as most edits resulted in skipping exon 53 or exon-reframing INDELS (i.e. 3n + 1) (**Fig. 4c**). To assess whether these genomic edits translate into a clinically relevant outcome, we measured the amount of dystrophin produced by the edited cells. We differentiated the myoblasts into myotubes for 10 days to enable production of dystrophin^51^. Using a dystrophin standard curve to quantify the restoration relative to a healthy myotube control, we observed dystrophin expression following treatment with the myo-sgRNA but not the liver-sgRNA (**Fig. 4d,e and Supplementary Fig. 7a**). We confirmed the production of dystrophin with immunofluorescence labelling (**Fig. 4f**). These results demonstrate that CRISPR MiRAGE can restore the production of dystrophin in DMD patient-derived myoblasts.

**Fig. 4:**
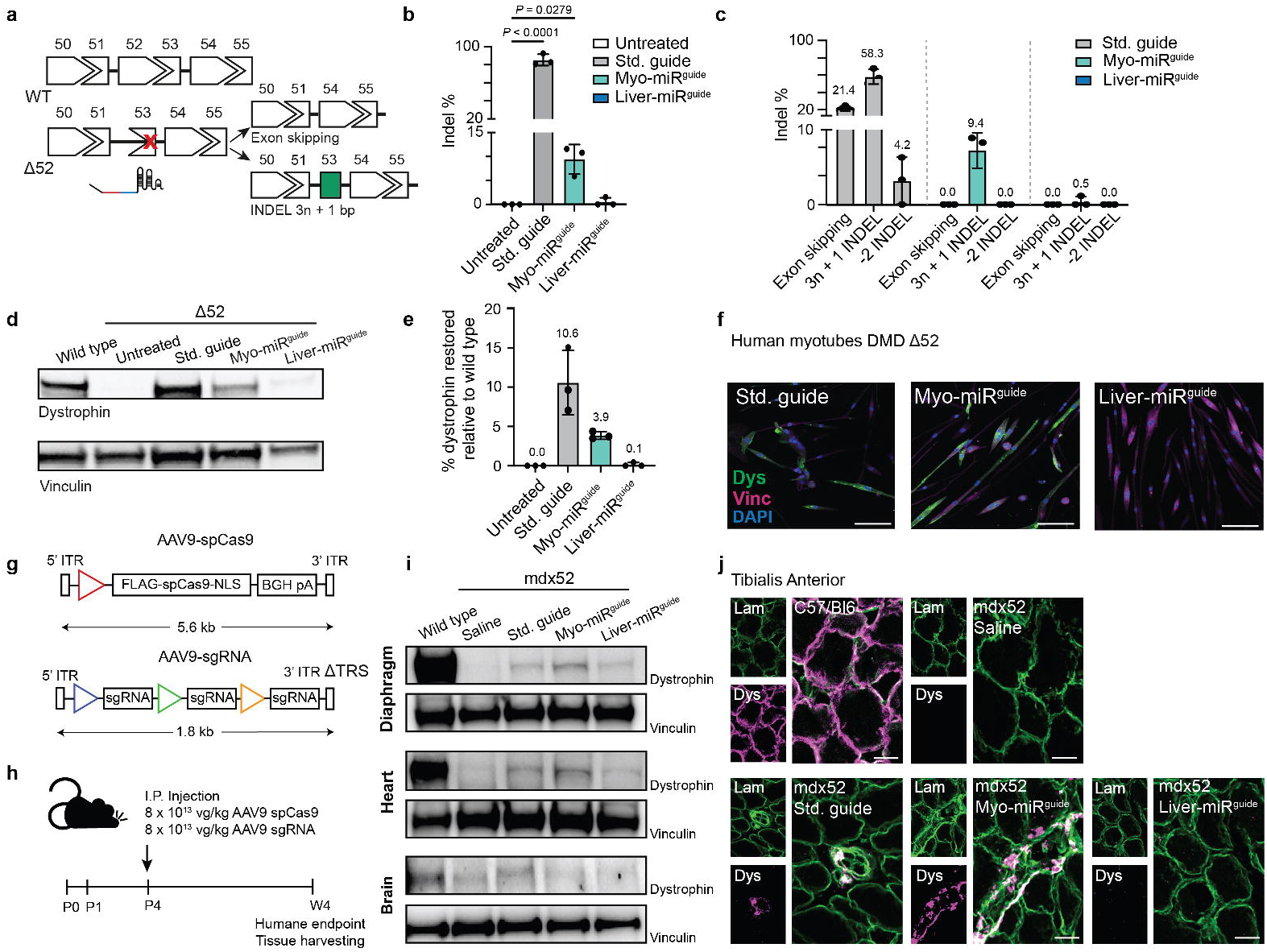
CRISPR-MiRAGE deployment in models of Duchenne Muscular Dystrophy. **a**. Diagram of the treatment strategy. Deletion of exon 52 (Δ52) in the *DMD* gene results in the formation of a premature stop codon in exon 53. Delivery of CRISPR MiRAGE using an optimized sgRNA targeting exon 53^46^, restores the correct reading frame of the *DMD* transcript by inducing skipping of exon 53 or reframing by precise insertion of 3n + 1 base pairs (single-cut strategy)^46^. **b**. Human Δ52 myoblasts were electroporated with a plasmid encoding SpCas9 and a sgRNA expression cassette. RNA retrotranscribed using *DMD*-specific primers was sequenced: % of INDEL events is shown (*n* = 3). **c**. Distribution of edited transcripts upon treatment with a standard sgRNA (std guide) and with miRNA sgRNAs responsive to muscle-specific miRNA (myo-miR-guide) and to liver-specific miRNA (liver-miR-guide) (*n* = 3). **d**. Representative western blot analysis of edited Δ52 human myoblasts differentiated into myotubes for 10 days. Samples were compared to a dystrophin standard curve using a protein lysate comprising different % of wild type human myotubes mixed with Δ52 human myotubes (*n* = 3). **e**. Quantification of dystrophin restoration for each replicate normalized to vinculin and relative to a wild type dystrophin standard curve (*n* = 3). **f**. Immunofluorescence staining of a representative set of edited human Δ52 myotubes (*n* = 1) showing Dystrophin (green), Vinculin (magenta) and nuclei (DAPI, blue). Scale bar 100 μm. **g**. Diagram depicting the constructs used within an AAV serotype 9 vector (AAV9). Due to the large size of SpCas9, we followed a dual-AAV strategy, one expressing SpCas9 under a strong ubiquitous promoter (Hybrid Chicken β-Actin, red triangle), and a second self-complementary AAV9 expressing the optimized sgRNA under 3 different Pol III promoters (U6, blue triangle; 7SK, green triangle; H1, yellow triangle). **h**. Mdx Δ52 mice at 4 days post-natal (P4) were injected intraperitoneally (I.P.) with 8 × 10^13^ viral genomes/kg of each virus. Four weeks later (W4), animals were sacrificed, and skeletal muscles and liver were collected. **i**. Representative western blots for diaphragm, heart, and brain. **j**. Immunofluorescence staining from Tibialis Anterior (TA) transversal tissue slices depicting dystrophin restoration upon treatments. Dystrophin is shown in magenta, Laminin in green, co-localization in white. Scale bar 20 μm. All data were analysed using one-way ANOVA and Dunnet’s multiple comparison test. The data represent the mean ± S.D.

We also tested our technology in a proof-of-principle study in the *mdx52* mouse model^52^. This animal model was chosen because it harbours the same genetic alteration in the *DMD* gene as in human myoblasts. However, a change of guide sequence is required to account for the murine origin of the mdx52 *DMD* gene^46^. Due to the packaging size limitation of our vector of choice, the recombinant adeno-associated virus serotype 9 (rAAV9), we generated two viruses, one expressing SpCas9 under the ubiquitous promoter CMV early enhancer/chicken β actin (CAG) promoter and the second expressing any one of the sgRNAs (positive control, myo-miR-guide or liver-miR-guide) in three copies, each under a different Pol III promoter (U6, H1 and 7SK), to increase its expression, as previously described^46^ (**Fig. 4g**). On post-natal day 4 (P4) a cohort of mdx52 male pups were randomised to receive intraperitoneally AAV9 SpCas9 and saline or any one of the three AAV9 sgRNAs (standard guide, myo-miR-guide or liver-miR-guide) (*n* = 3) (**Fig. 4h)**. A dose of 8 × 10^13^ viral genomes/kg was chosen for each AAV9, as previously established^53^. Four weeks post-injection (W4), the mice were sacrificed and tissues collected for analysis (**Fig. 4h**). We observed dystrophin protein restoration in diaphragm muscle and heart to a level comparable to the positive control upon deployment of the myo-miR-guide. Significantly lower levels of restoration were observed with the liver-miR-guide. On the other hand, no dystrophin restoration was detected in tissues not targetable with the myo-miR and liver-miR-guides, such as brain. However, we observed partial dystrophin restoration with the standard guide, overall supporting the specificity of CRISPR MiRAGE (**Fig. 4i, Supplementary Fig. 7b-d** and **Supplementary Fig. 8a-c**). We corroborated these results by performing immunofluorescence analysis in tibialis anterior (TA) muscles which showed dystrophin colocalized with laminin in mice treated with the myo-miR-guide and not with the liver-miR-guide (**Fig. 4j**). Together, these data show that rational sgRNA design enables control of SpCas9 editing activity based on tissue-specific miRNA activity in mammalian systems both *in vitro* and *in vivo*.

## DISCUSSION

With an increasing number of molecules approved for clinical use, nucleic acid-based therapies are rapidly emerging as a promising class of biotherapeutics capable of targeting the genetic bases of many human diseases. Currently, some of the biggest obstacles to their clinical translation are the risk of off-target activity, genotoxicity, and sub-optimal delivery, causing detrimental effects in cells and organs not directly affected by the disease. Our CRISPR-based gene editing approach addresses these limitations: the guide RNA becomes therapeutically active only upon interaction with specific, characteristic components of the target cellular environment, i.e. miRNAs. CRISPR MiRAGE relies on the introduction of dynamic secondary structure in the guide RNA that abolishes activity until disrupted by AGO-mediated miRNA recognition (**Fig. 1-3**). While a mechanism involving other ancillary RBPs cannot be excluded, simple strand displacement^54^ is not sufficient to induce the RNA conformational changes required to activate productive editing. The trigger hairpins contain targeted mismatches which are necessary to fine-tune their free energies and to control AGO interaction dynamics.

Although predictable folding has been reported to be a crucial limiting factor in the design of RNA devices^39,47,48^, we have shown that our miRNA-sensing sgRNAs fold into functional molecules regardless of the folding pathway followed (**Supplementary Fig. 4**). They perform as well when co-transcriptionally transcribed within the cell as when transfected following *in vitro* transcription and annealing (**Supplementary Fig. 6**). This provides flexibility to use a wide range of carriers, from adeno-associated viruses^49^, which require *in situ* transcription of the sgRNA, to lipid nanoparticles^44,45^, which require chemically synthesized sgRNAs^50^. In order to enhance CRISPR MiRAGE specificity and clinical applicability, given that a disease state is more likely characterized by a miRNA signature rather than a single miRNA, we also implemented a multi-sensing version, capable of responding to several predefined inputs, (**Fig. 3c**).

We have validated CRISPR MiRAGE in *in vitro* and *in vivo* models of DMD, a genetic disease mainly affecting skeletal and cardiac muscle, and showed that this tool is able to control SpCas9-mediated gene editing based on tissue-specific miRNA activity in a disease context (**Fig. 4**). The overall editing efficiency of CRISPR MiRAGE, as measured using the Stoplight reporter, is comparable to standard gene editing approaches that lack its capacity for context-sensitive control. However, the muscle-specific guide targeting the exon 53 of the *DMD* gene could be further improved. Previous evidence showing that sgRNAs targeting other DMD mutations achieve significantly higher editing levels^46^ and the observation that the genomic context of a target sequence plays a critical role on CRISPR activity^51^ suggest that optimization of the target guide sequence is necessary in order to advance such technologies towards clinical testing. Notably, although minimal, we observed dystrophin production also in skeletal muscles of mdx52 mice treated with the liver-miR-guide (**Fig. 4i**). There are previous reports showing that miR-122-5p, earlier described as liver-specific^25^, can be active in other tissues such as muscle, with high degree of variability across mouse strains and between human subjects^52^. We have also observed that levels of expression of miRNA do not necessarily correspond to miRNA activity (**Supplementary Fig. 1**). This suggests that, in order to accomplish the intended precise outcomes of this technology, thorough miRNA tissue profiling is needed and miRNA expression profiles should be coupled with tissue-specific miRNA activity estimates, for example using reporter systems or computational techniques^36^.

In conclusion, by rationally designing sgRNAs to sense and respond to tissue-specific miRNAs, we show here the potential of AGO-dependent RNA nanodevices for controlling the CRISPR gene editing in response to environmental cues. By addressing a fundamental hurdle of CRISPR editing, which is the risk of unintended editing events in bystander tissues, CRISPR MiRAGE holds great potential for enabling the use of such technologies to treat human diseases.

## MATERIALS AND METHODS

### Cell culture conditions

HEK293T Stoplight^+^ SpCas9^+^ were generated as previously described^34^. These cells were cultured in Dulbecco Modified Eagle’s medium (DMEM, Gibco) supplemented with 10% Fetal Bovine Serum (FBS, Gibco) and 1% Antimicotic-Antibiotic Solution (Sigma) at 37°C in a 5% CO_2_ atmosphere. Cell medium was changed every 48h, and cells were typically passaged at 90% confluency. We obtained human DMD Δ52 myoblasts (KM571) from Dr Vincent Mouly (Center for Myology, GH Pitié-Salpétrière, Paris, France). Human primary myoblasts were grown in Skeletal Muscle Cell Growth medium (PromoCell) supplemented with 1% Antibiotic/Antimycotic Solution (Sigma). When myoblasts were differentiated, Skeletal Muscle Differentiation Medium (PromoCell) supplemented with 1% Antibiotic/antimycotic (Sigma) was used. Myoblasts were left to differentiate for 10 days.

### Design of the miR-guides

Immediately upstream of a standard Cas9 guide sequence we introduced a miRNA binding site complementary to our miRNA of interest or to its seed sequence. To form the trigger hairpin, we added a shield sequence that is partially complementary to the miRNA binding site, the guide sequence, and the first G from the repeat sequence. In the case of a full-length miRNA-binding site, the partially complementary sequences reach no further than position 16, leaving the rest of the miRNA sequence to form the hairpin loop. For the miRNA seed design, the full seed is covered and a GAAA tetraloop was added. Nucleotides mismatched to the guide sequence were incorporated periodically (1-3 mismatches every 2-4 nucleotides). All Cas9 guide sequences incorporated 17-nucleotide targeting sequences, except for the dystrophin targeting sequence whose length was 20 nucleotides. Sequences of the sgRNAs are listed in Supplementary Table 1.

### sgRNA PCR assembly, plasmid construction, and purification

The sgRNA sequences were split into 2 parts, core and miRNA binding site, and ordered as separate oligonucleotides (Integrated DNA Technologies). The core consists of a reverse sgRNA backbone and a forward guide sequence with partial complementarity to the backbone. The core was PCR amplified in a 10 μL reaction with Q5 2X Mastermix (NEB) and primers at 500 nM concentration for 15 cycles using an annealing temperature suitable for the core sequence. The resulting PCR fragment was re-amplified in the presence of the microRNA binding site oligo, containing the miRNA binding site and a 25 nt overhang to the pU6 plasmid, and a reverse oligo, containing a 25 nt overhang to the expressing plasmid. A total of 0.5 μL from the core PCR reaction was added as a template to a 50 μL Q5 PCR reaction at 500 nM primer concentration for 40 cycles. The final product was run on a 15%, 29:1, 1 X TAE polyacrylamide gel. The gel was stained using SYBR gold (Invitrogen), and each sgRNA band cut out, submerged in 200 μL 0.5 M NaCl_2_, 1 X TE and left overnight. The following day, each band was spun down and the supernatant purified using a PCR clean up column (Qiagen). Each sgRNA was cloned into a U6 promoter-containing plasmid (modified from Hanson B. *et al*., 2022^53^). The sgRNA-expressing plasmids were built using the HiFi DNA Assembly Kit (New England Biolalbs), following the manufacturers protocol.

### Cell transfections

Cells were reverse transfected as follows. One hundred ng/well RNA was complexed with 0.2 μL Lipofectamine 2000 (Thermo Scientific) in the case of the miR-guides, or 0.1 μL Lipofectamine RNAiMAX in the case of the miRNA mimics and antagomirs, in 10uL of OptiMEM without antibiotics. Then, 20,000 HEK293T Stoplight^+^ SpCas9^+^ cells per well were plated in a 96 well plate (CytoONE, TC-treated, StarLab) in 100 μL complete DMEM and supplemented with 10 μL of plasmid-lipofectamine complexes. The mixture was left in the medium for 72h before eGFP/mCherry signal acquisition. miRNA 206 mimic and miR-17-5p antagomiR-were ordered from IDT and prepared as a stock at a suitable concentration. For experiments in human myoblasts, plasmid PX458 (Addgene plasmid 48138) was cloned to include sgRNAs with an optimized backbone and guide sequence hE53g10^46^. Control sgRNAs, a miR-206-3p and 122-5p-sensing sgRNAs were cloned into PX458. One μg of plasmid was electroporated into 200,000 human primary myoblasts carrying the Δ52 DMD mutation using the Neon® transfection system (Life Technologies, Invitrogen), following the manufacturer’s instructions. After electroporation, 1000 eGFP^+^ myoblasts cells were sorted using a BD Aria III cell sorter (BD Biosciences) and left growing as a polyclonal population. For the *in vitro* transcribed sgRNAs each sgRNA stock was made in 1X PBS and annealed (95 °C 3 min, −1 °C /3 sec until 20 °C) to promote proper sgRNA folding. Each sgRNA was used at a 50 nM final concentration per well, and cells were transfected as described above.

### eGFP/mCherry quantification assay

We use the ratio of fluorescence intensities from eGFP and the constitutively expressed mCherry as a measure of the level of activated editing for a given miRNA. Fluorescence was measured using a plate reader in well-scanning mode (Clariostar Plus, BMG Labtech), which allows the precise measurement of eGFP and mCherry fluorescence throughout the whole well surface. The cell culture medium was removed from the 96 well plate and the cells washed twice with 100 μL PBS supplemented with 0.8 mM MgCl_2_ and 0.9 mM CaCl_2_ to prevent cells from detaching. Then, 100 μL of divalent cation-supplemented PBS was added to each well, including 4-5 empty wells to be used as blanks. The well-scanning program uses a 10 × 10 measuring matrix in a 4 mm diameter for both eGFP and mCherry signals, with gain and focus set to automatic using the Enhanced Dynamic Range (EDR) feature.

### RNA/cDNA preparation and RT-qPCR

A total of 5 × 10^6^ HEK293T Stoplight^+^ SpCas9^+^ were plated in a 6 well plate. The next day, RNA was extracted using TriZol (Invitrogen) according to the manufacturer’s protocol. RNA was retrotranscribed using the High-Capacity cDNA Reverse Transcription Kit (Applied Biosystems). RNA and miRNA were quantified by qPCR using Power SYBR Green Master Mix (Life Technologies) or Taqman Universal PCR Mastermix (Applied Biosystems) respectively supplemented with transcript-specific probes (Supplementary Table 2 and Supplementary Table 3), following the manufacturer’s protocol. Analysis was performed on an Applied Biosystems StepOnePlus™ real-time PCR system (Life Technologies). We established Ct 35 as a threshold for no miRNA expression. 10ng RNA was used for miRNA quantification; 10, 20, 50 and 100 ng were used for the titration. 2 μL of cDNA mix prepared with 1 μg RNA in 100 uL were used for qPCR quantification of gene expression.

### miRNA sensor construction and assay

Dual luciferase assays were performed with a dual luciferase reporter plasmid containing a multiple cloning site (MCS) downstream of the firefly luciferase cassette (Promega). The MCS was digested using NheI and SalI, and the linearized plasmid was purified using a 1%, 1X TAE agarose gel followed by a gel extraction column (Qiagen). The linear plasmid was assembled into a miRNA sensor by adding a miRNA binding site in the MCS. The 2 oligos containing the miRNA binding site and 25-nucleotide overhangs corresponding to the vector backbone sequence were ordered from Integrated DNA Technologies, and annealed (98ºC for 30 sec., −3ºC/s until 20 °C in 1X TE buffer). The plasmid was assembled using the HiFi DNA Assembly kit (New England Biolabs). Bacterial culture and plasmid purification were performed as previously described^54^. Luciferase activity in transfected HEK293T Stoplight^+^ SpCas9^+^ cells was measured at 48h using a Dual-Glo Luciferase Assay System (Promega). Activity was calculated as the ratio of Firefly and Renilla luminescence intensities, normalised to untreated (dual-luciferase plasmid with no miRNA binding site), and presented as 1-(Firefly/Renilla) to represent miRNA activity as a positive number.

### NUPACK and CoFold

We used the NUPACK web server (https://www.NUPACK.org)^33^ and submitted all RNA samples for analysis at 37 °C with a maximum complex size of 1. Free energy parameters used were those determined by Serra and Turner^34^ with “some” in the dangle tab. Salt conditions were 1 M NaCl. No pseudoknots were allowed. For structural predictions accounting for co-transcriptional folding, we used CoFold (https://e-rna.org/cofold/)^38^. Structures for all miRNA-sensing sgRNAs were calculated using the thermodynamic energy parameters stated in Turner^55^ with default scaling parameters (α = 0.5 and τ = 640).

### RNA *in vitro* transcription and purification

T7 promoter-containing sgRNA templates were PCR-amplified using Q5 master mix and gel-purified with a polyacrylamide gel. Purified templates were used in 20 μL T7 *in vitro* transcription reactions using the HiScribe T7 *in vitro* transcription kit (New England Biolabs) at 100 nM, and left to react at 37 °C overnight. Next morning, each reaction was topped up to 100 μL, supplemented with 1 X CutSmart buffer (New England Biolabs) and 6 μL Quick CIP (New England Biolabs), and left to react for 3 h at 37 °C. sgRNAs were cleaned up using Trizol as described by the manufacturer. The clean RNA was gel-purified using an 8.5 M Urea, 15%, 29:1, 1X TAE polyacrylamide gel alongside an RNA ladder (Riboruler, Thermo Scientific); each sgRNA was loaded using 2X RNA loading dye (New England Biolabs). sgRNAs were heated for 3 min at 95 °C prior to loading into the gel. The gel was stained with SYBR gold and the bands corresponding to the full-length sgRNA were cut out, submerged in 0.5M NaCl 1X TE overnight and the supernatant subjected to Trizol extraction.

### Flow cytometry

60,000 HEK293T StopLight^+^SpCas9^+^ were plated in a 24 well plate and reverse transfected with 5 ng miR-18a-5p guides with optimized backbone and co-transfected with miR-18a-5p antagomiRs at a final concentration of 500 nM where indicated. Cells were collected 72 h later for cytometry analysis. All conditions were washed in PBS and the resulting cell pellets were resuspended in 250uL PBS supplemented with 2% FBS. Cells were kept on ice until sample acquisition. We acquired a minimum of 50,000 gated-cells per sample. Spectral flow cytometry data was acquired on a spectral ID7000 cytometer (Sony Biotechnology). All the spectral data were unmixed using the WLSM algorithm of ID7000 software. Then, the unmixed sample data were converted to FCS files and analysed using FlowJo version 10.9 software (Treestar). Unmixing matrices were set using cells and compensation beads (BD Biosciences). Gating strategy shown in **Supplementary Fig. 6e**.

### Human primary myotubes RNA extraction and transcript analysis for INDELS

Human primary myoblasts, electroporated with the sgRNA and sorted as described above, were differentiated for 10 days, changing the medium every 48 h. On day 10, cells were harvested and their RNA was extracted using the Qiagen RNEasy Kit, following the manufacturer’s instructions. cDNA was prepared using the Superscript III One-step RT-PCR system with Platinum Taq (Thermo Scientific) using 200 ng of RNA and DMD specific primers at a final concentration of 200 nM in a 25 μL RT-PCR reaction. The cycling conditions were as follows: 30 min at 60 °C, 2 min at 94 °C, 35 cycles with 15 s at 94 °C, then 60 sec at 60 °C, and 90 sec at 68 °C, and a final cycle for 5 min at 68 °C. Then, a nested PCR was performed to the RT-PCR product using AmpliTaq Gold DNA polymerase (Thermo Scientific). Each 25 μL Nested PCR reaction was prepared with 1.25 μL from each RT-PCR mix as template, using DMD-specific primers at a final concentration of 200 nM (Supplementary Table 3). The nested PCR used the following cycling conditions: 10 min at 95 °C, followed by 30 cycles with 40 sec at 94 °C, 40 s at 60 °C, and 60 sec at 72 °C, and a final extension cycle of 7 min at 72 °C. The PCR product was purified using a Qiagen PCR purification kit and sent for Sanger sequencing at an amplicon concentration of 10 ng/μL. Samples were sequenced using a DMD exon 54 reverse primer. Sanger sequencing results were deconvoluted using the DECODR online suite^56^, using the untreated Δ52 DMD myotube samples as control. Each condition was tested in triplicate.

### Immunofluorescence

Human myoblasts were grown in a 6-well plate containing a sterile coverslip. Once confluency was reached, the medium was changed to differentiation medium for 10 days. The medium was exchanged every 48 h. Human myotubes were fixed with 4% Paraformaldehyde diluted in PBS for 20 min. Each well was washed 3 times in PBS for 15 min. Myotubes were then blocked in 5% Normal Goat Serum (NGS, Gibco) and incubated with primary antibodies against dystrophin (15277, Abcam) and vinculin (V9131, Sigma) overnight at 4 °C. The next morning, cells were washed 3 times for 5 min with PBS before incubating with a secondary antibody (ab150080 and ab150157, Abcam) for 1 h at room temperature. Cells were washed a second time, dab dried, and mounted on a slide for imaging in an Olympus Fluoview FV1000 confocal microscope. Frozen tissue mounted on a cork was cut into 8 μm slices using a Cryostat. Tissue slices were blocked for 2 h at room temperature with 20% Fetal Bovine Serum (FBS, Gibco) and 20% Normal Goat Serum (NGS, Gibco), and left overnight at 4 °C with the primary antibodies against dystrophin (15277, Abcam) and α-laminin (L0663, Sigma) in 20% NGS. The next day, the slides were washed 4 times with PBS and stained with the secondary antibodies (ab150080 and ab150113, Abcam) for 1 h at room temperature in PBS. Slides were washed a second time and stained with Hoechst 33342 at 5 μg/mL (Thermo Scientific). After that, the slides were mounted and left 12-24 h for setting at 4 °C. Slides were imaged with a Zeiss 980 IDRM Airyscan 2 confocal microscope.

### Adeno associated virus design and production, and animal experimental design

AAV constructs were designed and submitted to VectorBuilder for cloning and packaging into an AAV9 vector. AAVs expressing the following transgene were produced: SpCas9, single-cut *dmd* targeting standard sgRNA (std sgRNA), a miR-206-3p-sensing sgRNA (myo-miR-guide) and a miR-122-5p-sensing sgRNA (liver-miR-guide). All sgRNAs contain guide sequence mE53g2^46^.

### Animals

The experimental design for the animal experiments was based on the three R principle (replacement, reduction and refinement) to minimize both suffering and the number of animals used. All procedures were approved by the Animal Investigation Committee of the National Institute of Neuroscience and the National Centre of Neurology and Psychiatry (Japan) (Approval number: 2019012). *Dmd* exon 52-deficient muscular dystrophy mice (mdx52 mice) have been backcrossed to the C57BL/6J (WT) strain for more than eight generations. The mice were allowed ad libitum access to food and drinking water. *Mdx52* postnatal 4 (P4) pups were injected intraperitoneally with 8 × 10^13^ viral genomes/kg of each virus in a single dose. The animals were sacrificed 4 weeks after treatment and the tissues were prepared for western blot and immunofluorescence.

### Tissue protein extraction and western blot

A piece of tissue was cut while frozen and resuspended in lysis buffer (Tris-HCl pH 6.8, 2% SDS, 15% glycerol) supplemented with a proteinase inhibitor cocktail (cOmplete mini-no EDTA, Roche). The tissue and buffer were moved to a pre-chilled master tube with Zirconia Beads (Zirconia Beads 3.0 mm, Biomedical Sciences) and disrupted using a Shakeman device (Biomedical sciences). The homogenised tissues were spun down briefly to remove bubbles and transferred to a heat block at 95 °C for 5 min. After heating, the sample was spun down for 15 min at 15,000*g* at 4 °C The supernatant was collected and used for protein quantification using a BCA assay (Thermo Scientific). Protein from myotubes was extracted using 1 X radioimmunoprecipitation assay (RIPA) lysis buffer supplemented with 10% SDS, and 1 X cOmplete protease inhibitor cocktail. For western blot we used 15 μg of protein lysate for myotube, 60 μg for heart and diaphragm, and 80 μg for brain. Protein lysates were mixed in NuPAGE™ LDS Sample Buffer (4X) (Thermo Scientific) and NuPAGE™ Sample Reducing Agent (10X) (Thermo Scientific) and heated at 70 °C for 10 min. The samples were loaded into a NuPAGE™ 3 to 8%, Tris-Acetate 1 mm gel (Thermo Scientific) with HiMark™ Unstained Protein Standard (Thermo Scientific) and run at 150 V for 90 min. The gel was then transferred to a PVDF membrane using 1X NuPAGE™ Transfer Buffer supplemented with 10% methanol and 0.1 g SDS/L. The transfer was run for 1 h at 30 V and another 1 h at 100 V. Once the transfer was finished, the membrane was stained with 1X Fast Green FCF (Sigma) for 15 min and imaged for total protein. Then, the membrane was blocked with Intercept® (PBS) Blocking Buffer (Li-Cor) for 1 h at room temperature. After that, the blot was stained with primary antibodies against dystrophin (NCL-DYS1, Leica), vinculin (V9131, Sigma) in blocking solution (LI-COR) with 0.1% Tween 20 overnight at 4 °C. The next day, the blots were washed with PBS supplemented with 0.1% Tween 20 four times for 5 min before staining with the secondary antibody (7076, Cell Signaling; ab216773, Abcam) in blocking buffer with 0.1% Tween 20 for 1h at room temperature. After the secondary antibody incubation, the blots were washed again and the HRP signal was developed using SuperSignal™ West Femto Maximum Sensitivity Substrate in a Bio-Rad ChemiDoc MP.

### Statistical analysis

Data were plotted using GraphPad Prism 9. All experiments were first submitted to a ROUT test (Q = 1%) to detect outliers. The resulting data were tested using a one-way ANOVA with a Dunnett’s or Tukey’s test for multiple comparison correction, as indicated. *P* value < 0.05 was set as significant. All experimental replicates are comprised of independent measurements.

## Data availability

The main data supporting the findings of this study are available within this paper and its Supplementary Information. The raw and analysed datasets are too large to readily share publicly yet are available for research purposes from the corresponding author on reasonable request.

## Acknowledgements

This study was supported by a Career Development Fellowship grant from the Medical Research Council to C.R. (MR/Y009703/1) and research grants from the Kennedy’s Disease Association to C.R. and from the Muscular Dystrophy Association/American Association of Neuromuscular & Electrodiagnostic Medicine Foundation (578222) to C.R. W.F.L. was supported by the Medical Research Council programme grant to M.J.A.W. (MR/N024850/1). A.J.T and A.G.G were supported by Marie Curie Initial Training Network EScoDNA [FP7 project 317110], and Oxford University Press John Fell Fund. A.G.G was also supported by a La Caixa Foundation Fellowship (ID: LCF/BQ/EU16/11560044).

## Author contributions

A.G.G., C.S., W.F.L., and J.R. conducted experiments and analysed the data. O.G.J., P.V., S.E.A., and J.B. provided materials and assistance. A.G.G. and C.R. wrote the paper, with input from all authors. A.G.G., Y.A., A.J.T., M.J.W., and C.R. supervised the study.

## Notes

### Competing Interest Statement

The authors have declared no competing interest.

### Summary of Updates

Figure 4 revised; author list revised.

