## Supplementary Figures for "Tissue-specific modulation of CRISPR activity by miRNA-sensing guide RNAs"

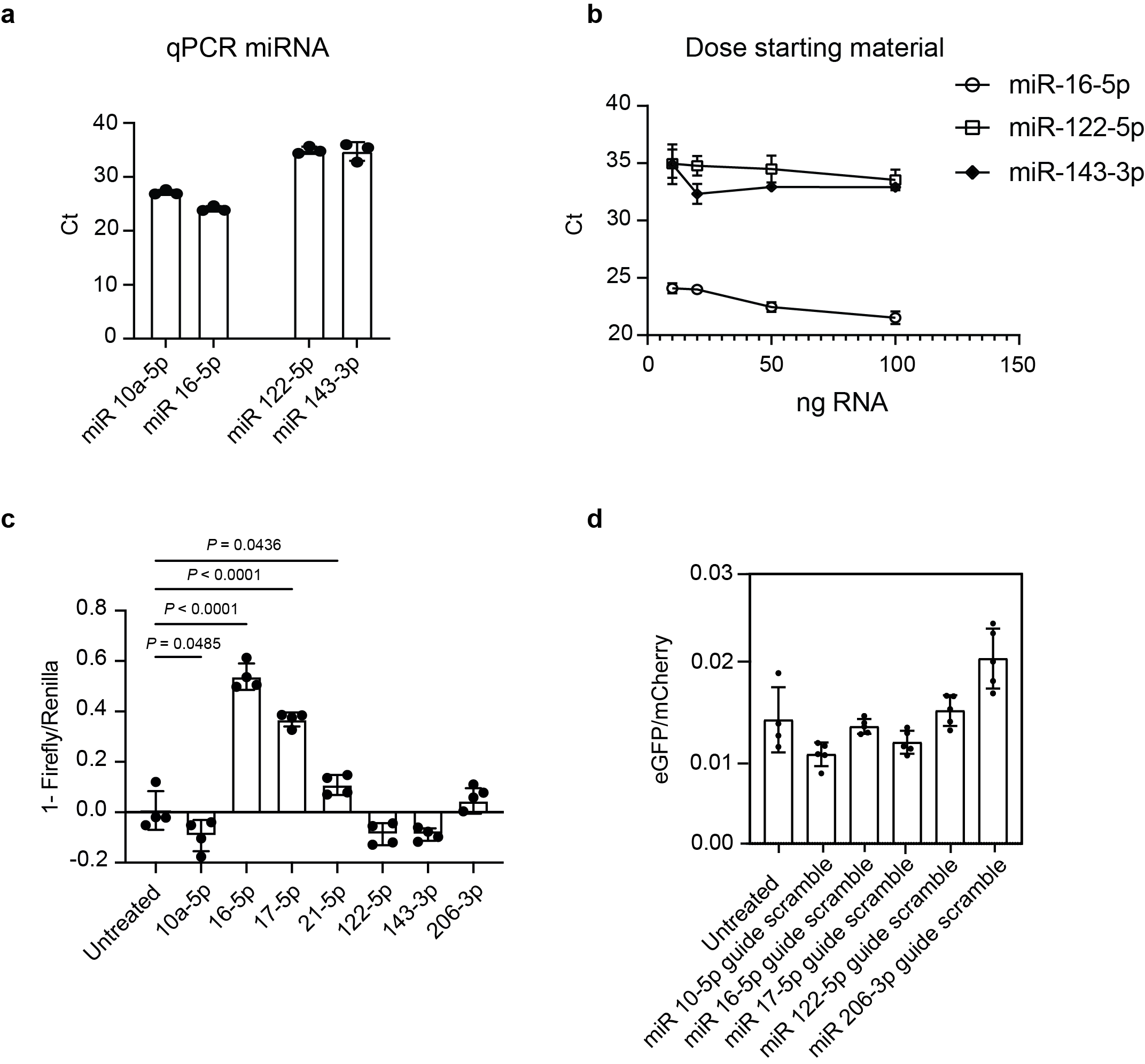

**Supplementary Figure 1: a.** Expression of miR 10-5p, miR 16-5p, miR 122-5p and miR 143-3p in HEK293T Stoplight^+^spCas9^+^ assayed by qPCR. The threshold for detection of expression was set at Ct = 35 cycles (*n* = 3). **b.** Dose curve using increasing amounts of starting material (extracted RNA) as template for cDNA production (10, 20, 50 and 100 ng of RNA). Expressed miRNA miR 16-5p shows a reduction in Ct values as the dose of starting material increases, whereas the Ct value for non-expressed miRNAs (miR 122-5p and miR 143-3p) remains approximately constant (*n* = 3). **c.** Dual-luciferase miRNA activity data. Sensors for 7 different miRNAs (miR 10a-5p, miR 16-5p, miR 17-5p, miR 21-5p, miR 122-5p, miR 143-3p and miR 206-3p) were built and transfected into the HEK293T Stoplight^+^spCas9^+^. Data was collected 48 h later and presented as 1-(Firefly/Renilla luciferase luminescence) to represent miRNA activity (*n* = 4). **d.** Activities of control miR sgRNAs with scrambled miRNA binding site sequences (cf. Figure 1e) (*n* = 5). All data were analysed using one-way ANOVA and Dunnet’s multiple comparison test. The data represents the mean ± S.D.

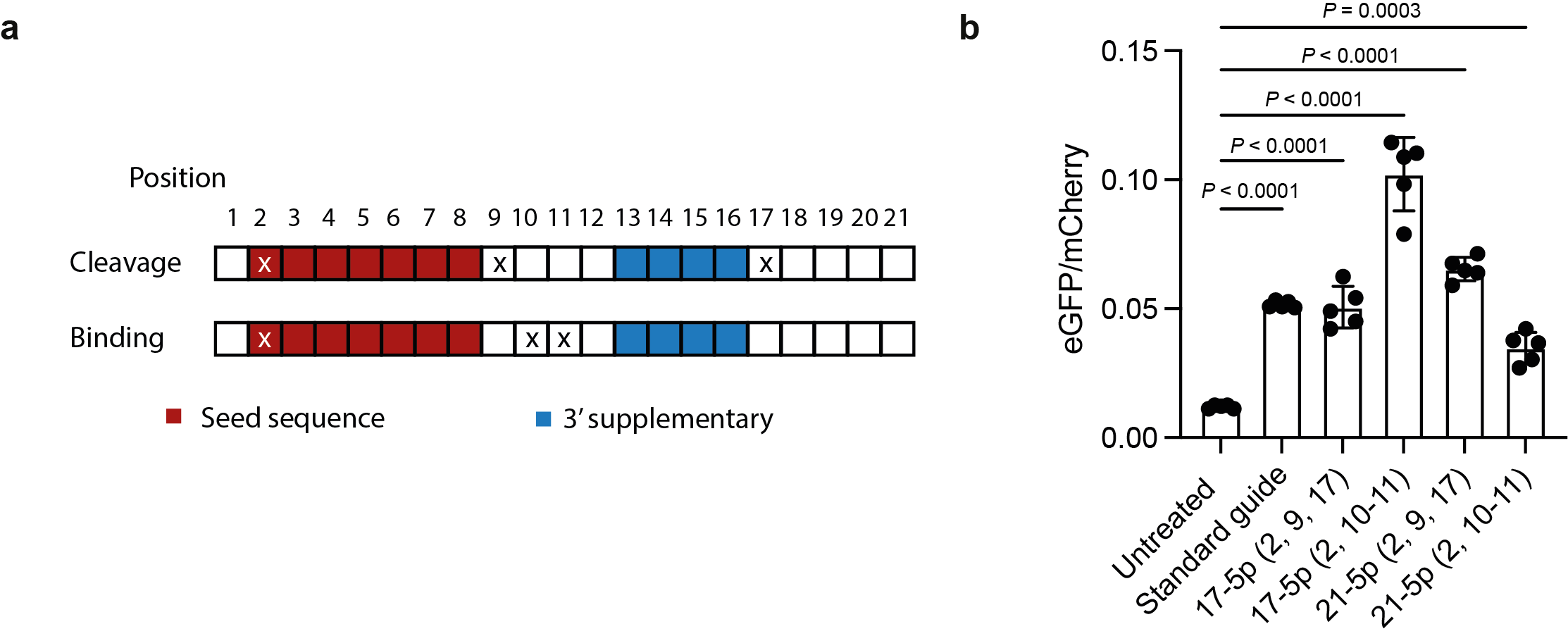

**Supplementary Figure 2: a.** Diagram depicting the location of the targeted mismatches based on their effect on RISC downstream modality^24^. **b.** RISC-mediated cleavage or binding as the miR guide activation mechanism. miR guides responsive to miR 17-5p and miR 21-5p were made with cleavage-permissive mismatches (2, 9, and 17) and cleavage-abolishing mismatches (2, 10-11). There are no significant differences between mismatch profiles suggesting that binding alone is sufficient to cause activation (*n* = 5). All data were analysed using one-way ANOVA and Dunnet’s multiple comparison test. The data represents the mean ± S.D.

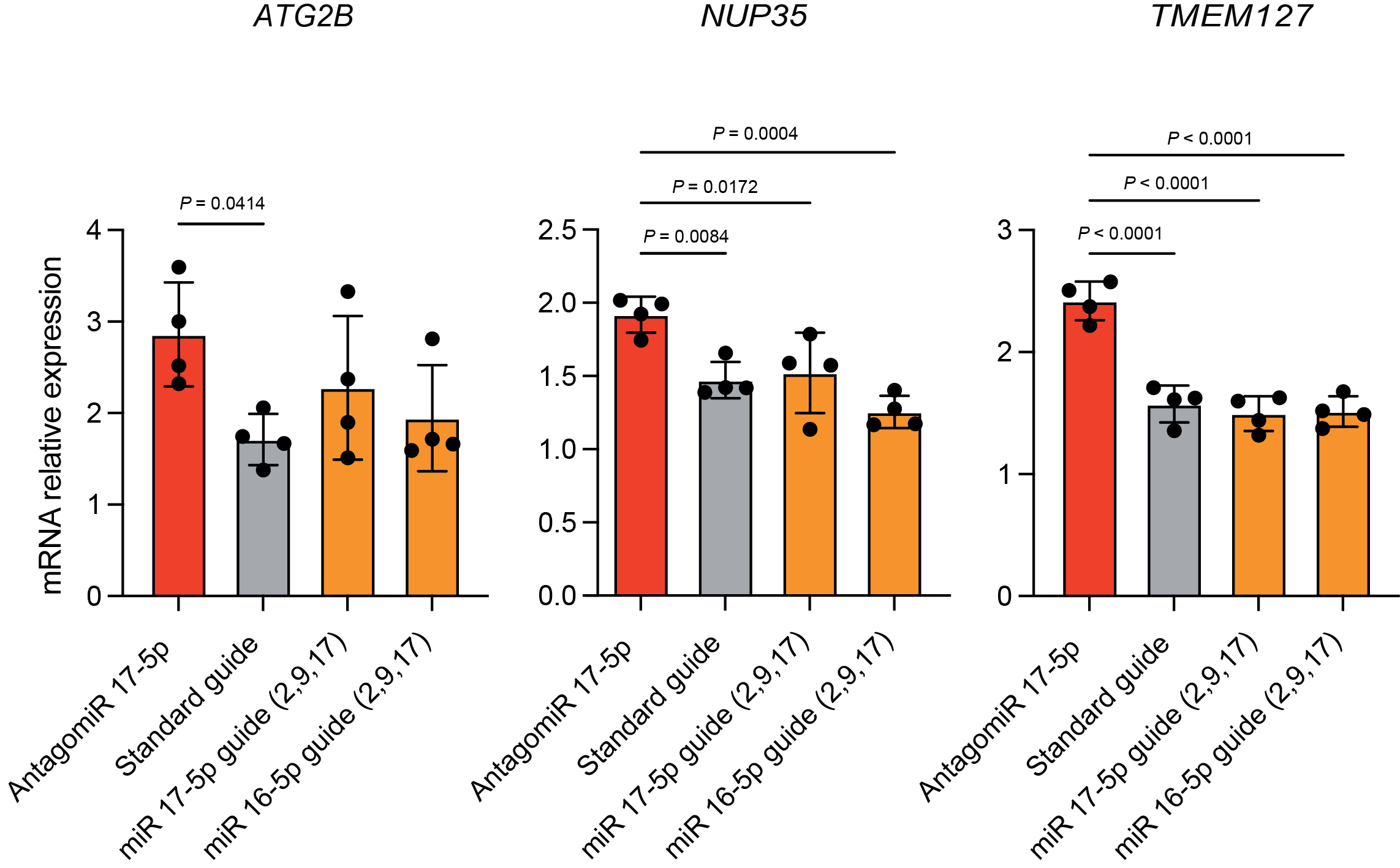

**Supplementary Figure 3:** Effect of miR 17-5p sgRNA on the expression of miR 17-5p-regulated transcripts (*ATG2B*, *NUP35*, and *TMEM127*) relative to *HPRT* housekeeping gene and normalised to an untreated control. While treatment with the miR 17-5p antagomir resulted in increased expression of the indicated targets, no statistically different changes in expression were observed between miR 17-5p sgRNA, a standard sgRNA (standard guide) and a miR 16-5p sgRNA, suggesting that the trigger hairpin does not affect mRNA homeostasis (*n* = 4). All data were analysed using one-way ANOVA and Dunnet’s multiple comparison test. The data represent the mean ± S.D.

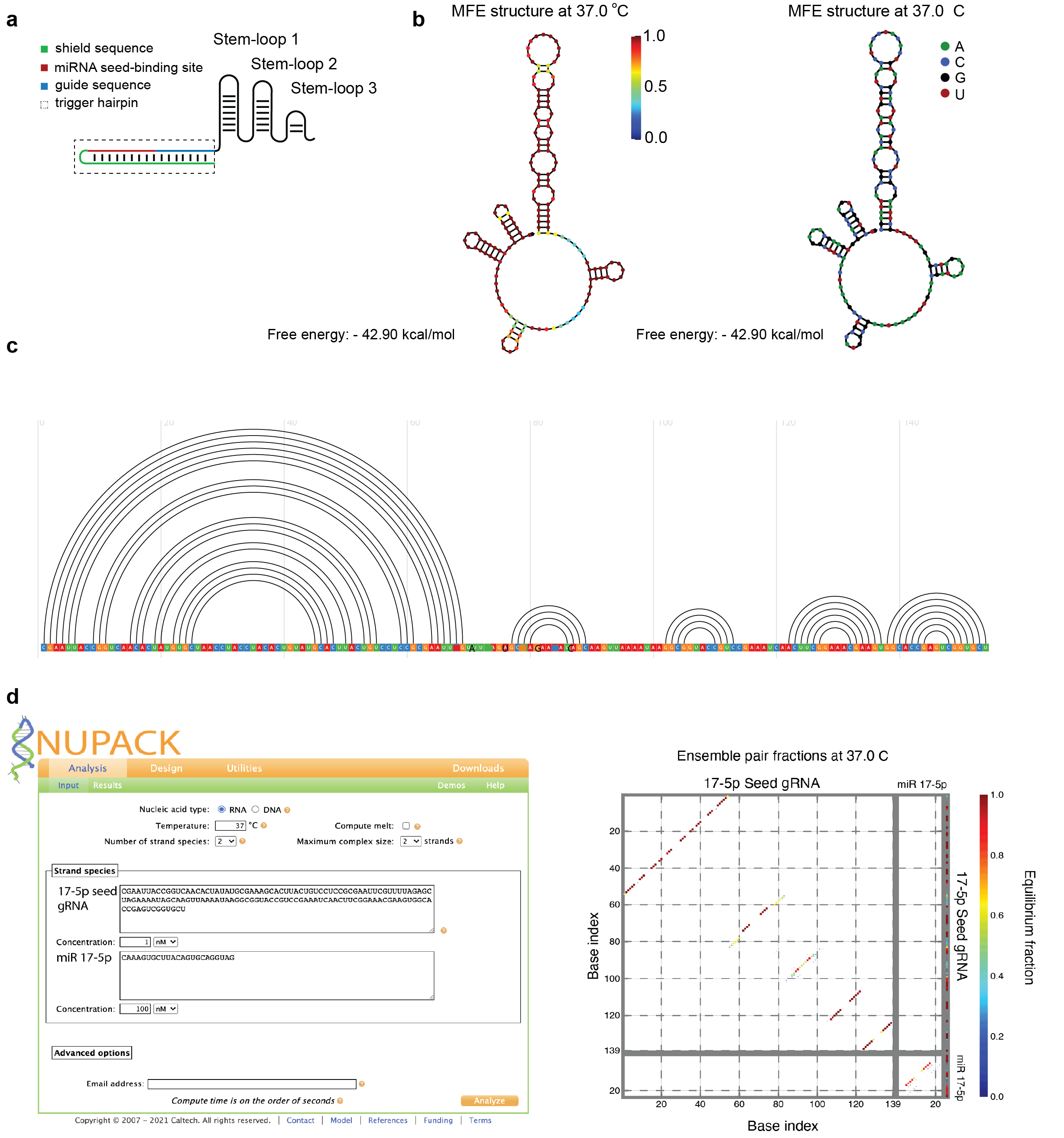

**Supplementary Figure 4:** Structure of a miR guide. **a.**
Structural features of a miR guide. **b** Using NUPACK^32,33^, we calculated the most common conformation in the equilibrium ensemble of the miR guide. The probability that each nucleotide is found in the specified position (left) and a colour-coded depiction of the miR guide sequence in this structure (right) are shown. **C.** CoFold^37,38^ calculation of the secondary structure of the same miR 17-5p miR guide after simulated co-transcriptional folding. **d.** Equilibrium secondary structure of miR 17-5p guide (1 nM) in the presence of miRNA 17-5p (100 nM) calculated using NUPACK. Left panel shows the parameters, species, sequences and concentrations used. We chose a 100 times excess miRNA to sgRNA ratio to favour hybridization, while the concentration was set to the nM range based on typical miRNA concentrations in a mammalian cell. Right panel shows the hybridization map indicating the equilibrium probability of each possible hybridized base pair. No inter-strand interaction is predicted, suggesting that the mechanism of activation of the miR guide is not simple strand displacement and is therefore likely to be mediated by AGO, as designed.

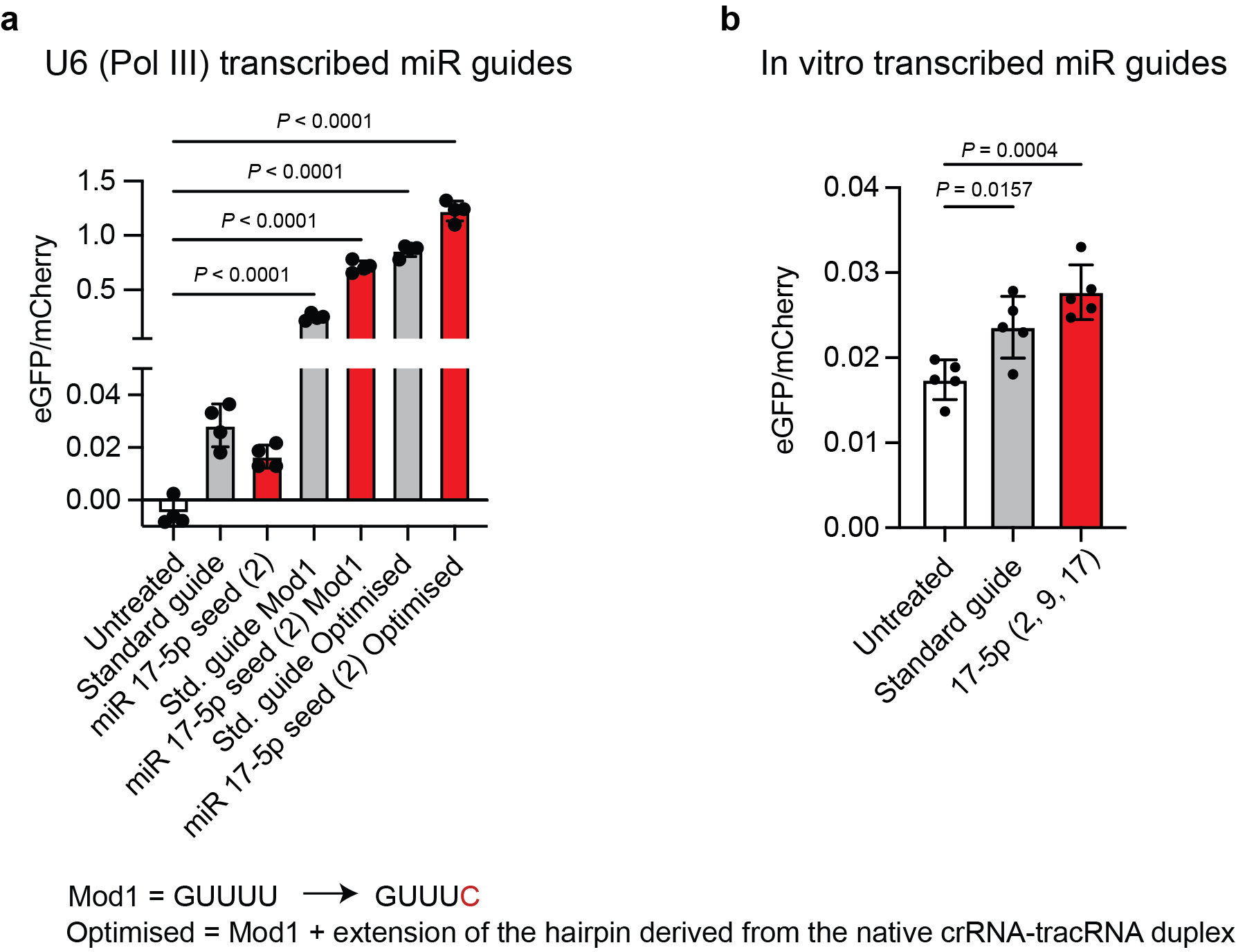

**Supplementary Figure 5:** sgRNA backbone optimisation increases activity. **a.** The activities of miR guides transcribed from a Pol III promoter increase by approx. 50× when the four consecutive Us in the repeat region are changed to three Us and one C (labelled as Mod1), reducing polymerase stalling^46^. (Note the change in axis scaling at the break). A further known modification of the sgRNA backbone, extension of the hairpin derived from the native crRNA-tracRNA duplex^46^, resulted in a further increase in activity to approx. 75× (labelled as Optimised). (*n* = 4). **b.** Transfection of an *in vitro* transcribed active (miR 17-5p) at a final concentration of 50 nM resulted in detectable editing activity (*n* = 4). Data were analysed using one-way ANOVA and Dunnet’s multiple comparison test. The data represent the mean and the error bars, SD.

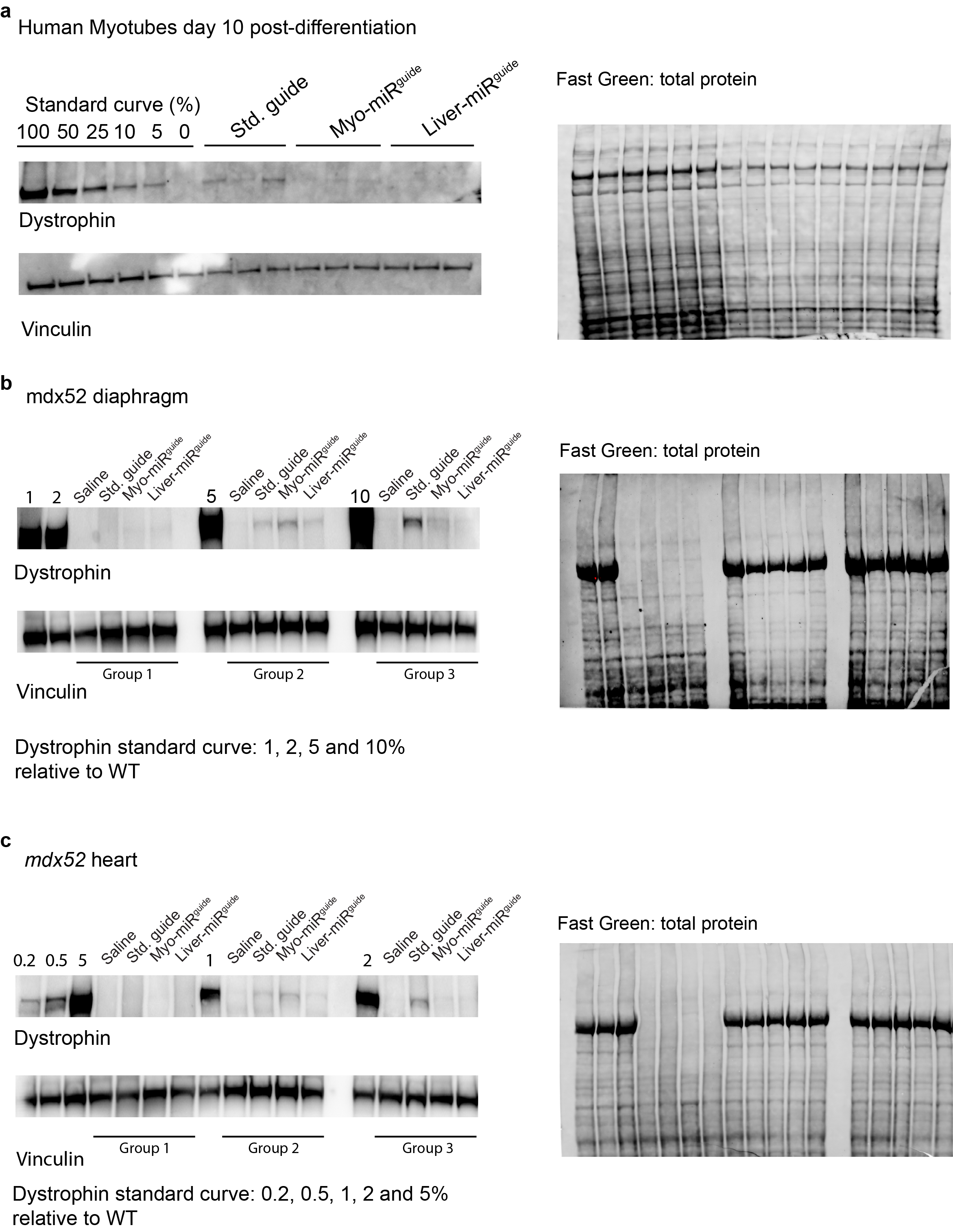

**Supplementary Figure 6:** Quantification of dystrophin restoration. **a.** Western blot used for the quantification of dystrophin restoration in Δ52 human myotubes shown in Figure 4e. A total of 15 μg of protein were blotted against dystrophin and vinculin (loading control). Total protein stain of the membrane using Fast Green FCF is shown on the right (*n* = 3). **b, c.** Western blots used for the quantification of dystrophin restoration in *mdx52* diaphragm and heart shown in Figure 4i. A total of 60 μg of protein were blotted against dystrophin and vinculin (loading control). Total protein stain of each membrane using Fast Green FCF is shown on the right (*n* = 3).

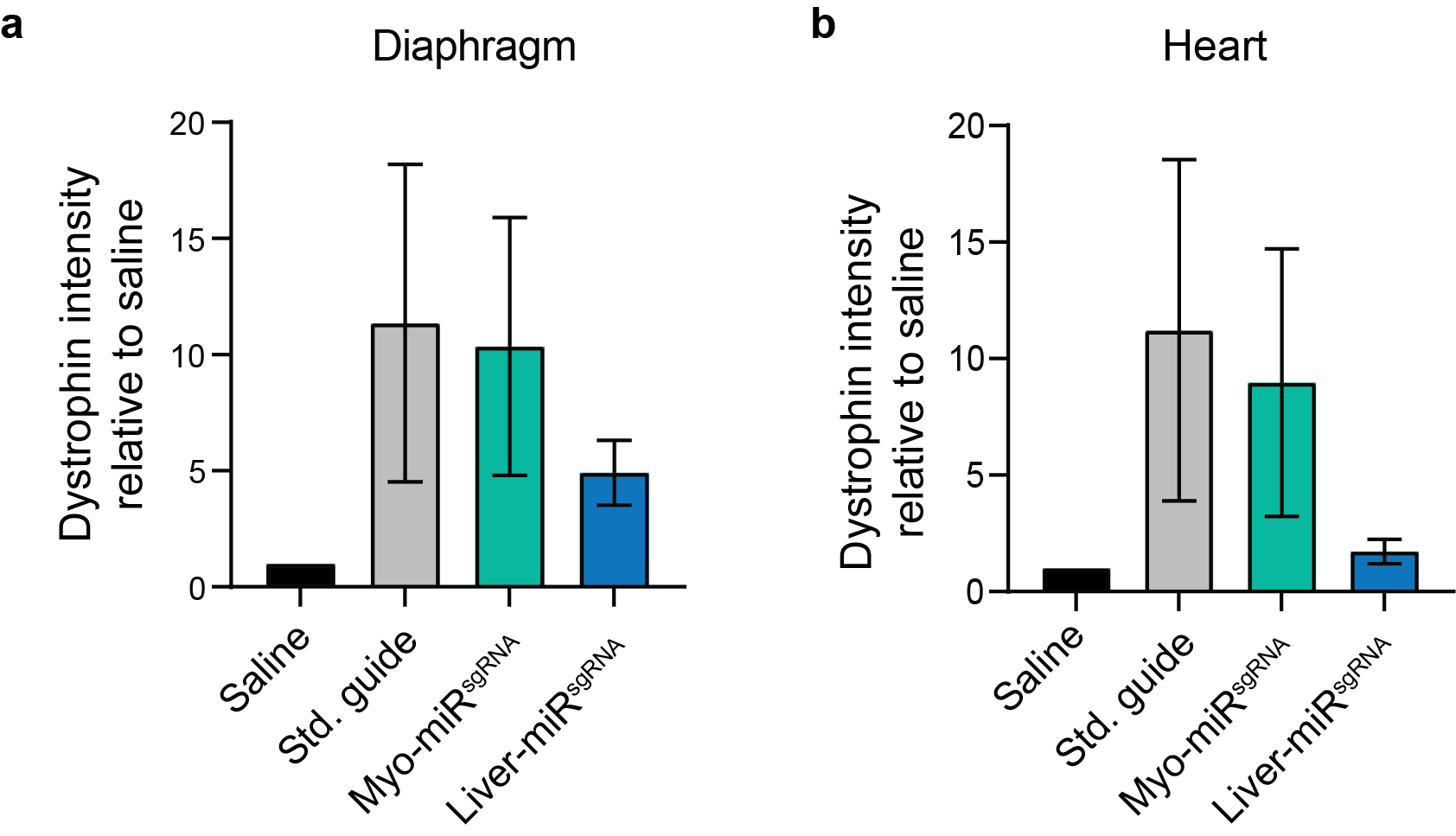

**Supplementary Figure 7:** Quantification of dystrophin production from the western blots in Supplementary Fig. 5. Dystrophin levels are normalized to those measured for saline-treated animals (*n =* 3). **a.** Diaphragm. **b.** Heart. The data represent the mean and the error bars the SEM.
